# Outlier Detection in Single-Cell Transcriptomics Reveals Disease-Enriched Cytotoxic Immune Populations

**DOI:** 10.64898/2026.01.12.699092

**Authors:** Devanshi Gupta, Arsh Gupta

## Abstract

Single-cell RNA sequencing (scRNA-seq) enables detailed analysis of cellular heterogeneity, but standard clustering approaches can miss rare cell populations with important disease relevance. We present an outlier-aware framework that complements clustering by identifying statistically unusual cells. Applied to COVID-19 bronchoalveolar lavage fluid (BALF) samples, this approach revealed strong disease-associated enrichment of outliers: 9.62% in moderate disease and 6.16% in severe disease, compared with 1.14% in healthy controls. These cells exhibited elevated cytotoxic marker expression (NKG7, GZMB, PRF1, GNLY; 2.0–2.9× enrichment) consistent with activated effector immune states, with replication across independent datasets. To verify that outliers were not low-quality cells, we confirmed superior QC metrics: outliers showed 68% more genes and 78% higher UMI counts than normal cells. Application to rheumatoid arthritis PBMCs showed modest enrichment (5.40% in RA vs. 4.66% in healthy), consistent with chronic inflammation. Comparison of four outlier detection algorithms identified Isolation Forest as the best-performing method. Subclustering revealed diverse functional states within outliers, including NK cytotoxic cells, activated T cells, and proliferating cells. Overall, outlier detection reveals rare, functionally distinct immune populations overlooked by conventional clustering.

## 1 Introduction

Single-cell RNA sequencing (scRNA-seq) has transformed our ability to study cellular heterogeneity, revealing rare cell populations invisible to bulk measurements. However, the dominant analytical paradigm - unsupervised clustering - assumes cells organize into discrete, stable populations. This assumption breaks down for rare, highly activated cells that constitute less than 5% of samples but may disproportionately drive disease pathology.

Rare cell populations - typically less than 5% - can play outsized roles in disease. Examples include tissue-resident memory T cells in chronic inflammation, regulatory T cells in autoimmunity, and activated natural killer (NK) cells in viral infections. Standard clustering may force these cells into inappropriate clusters or discard them as low-quality outliers.

Outlier detection, a well-established machine learning approach, offers a complementary strategy. Rather than grouping all cells, outlier detection identifies observations that deviate significantly from the majority. Methods such as Isolation Forest, Local Outlier Factor, and One-Class SVM have been used successfully in fraud detection and anomaly monitoring, but are underutilized in single-cell transcriptomics.

COVID-19 presents a compelling context for studying rare immune populations. Severe disease is characterized by immune dysregulation and tissue damage, but the role of highly activated rare cells remains incompletely understood. RA, by contrast, represents chronic low-grade inflammation driven by dysregulated immunity.

Here, we introduce a systematic framework to identify and characterize outlier cells in scRNA-seq data. We applied this to COVID-19 (acute viral infection) and RA (chronic autoimmune disease), analyzing approximately 194,000 cells across three independent datasets.

## 2 Materials and Methods

### 2.1 Data Acquisition

#### COVID-19 Discovery

We obtained scRNA-seq data from BALF of COVID-19 patients from GEO (GSE145926): 3 healthy controls, 3 moderate patients, 6 severe patients (108,230 cells, 10 × Genomics). **Validation:** GSE147143 with 3 severe patients (118,421 cells). **Rheumatoid Arthritis:** PBMC data from 18 RA patients and 18 healthy controls from CellxGene (108,717 cells).

### 2.2 Processing

Raw count matrices were processed using Scanpy v1.9.3. Quality control retained cells with 200–6,000 genes, 500–40,000 UMI counts, and *<*20% mitochondrial expression. Data were normalized using counts-per-million and logtransformed. Highly variable genes (n=2,000) were identified, and PCA was performed (50 components). Leiden clustering (resolution 1.0, k=15) and UMAP were computed for visualization.

### 2.3 Outlier Detection

Outlier detection used Isolation Forest (scikit-learn v1.3.0) on 50-dimensional PCA space (n_estimators=100, contamination=‘auto’, random_state=42). We compared four methods: Isolation Forest, Local Outlier Factor (n_neighbors=20), One-Class SVM (kernel=‘rbf’, nu=0.05), and DBSCAN (eps=0.5, min_samples=5). Statistical significance was assessed using chi-square and Fisher’s exact tests. Gene expression comparisons used Wilcoxon rank-sum tests with Benjamini-Hochberg FDR correction. Outlier subclustering used Leiden (resolution=0.8), and marker genes were identified with log2 fold-change *>*0.5 and FDR *<*0.01.

### 2.4 Quality Control Validation

To verify that outliers were not low-quality cells, we compared QC metrics (number of genes, UMI counts, mitochondrial percentage) between outlier and normal cells using Mann-Whitney U tests.

## 3 Results

### 3.1 Outlier Enrichment in COVID-19

Applying Isolation Forest to COVID-19 BALF data (84,567 cells) identified outlier immune cells strongly enriched in disease (Fig. 1). Outlier frequency was 9.62% in moderate patients (825/8,579), 6.16% in severe (3,114/50,569), and 1.14% in healthy controls (290/25,419), representing 8.43-fold and 5.40-fold enrichment (*χ*^2^ = 1324, *p* <10^−288^).

**Figure 1:**
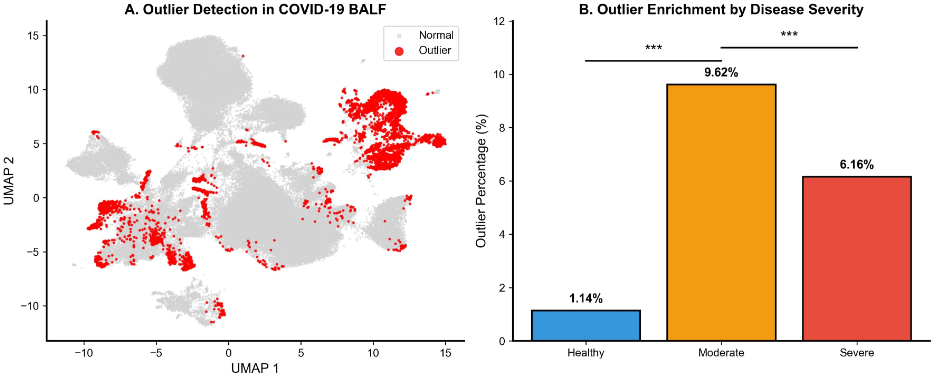
COVID-19 outlier detection and enrichment. (A) UMAP visualization of 84,567 cells from COVID-19 discovery cohort showing outlier cells (red) dispersed across the transcriptional landscape compared to normal cells (gray). (B) Outlier percentage by disease severity demonstrating significant enrichment in moderate COVID-19 (9.62%) and severe COVID-19 (6.16%) compared to healthy controls (1.14%). \*\*\**p* < 0.001 by Fisher’s exact test.

The higher frequency in moderate versus severe disease suggests peak immune activation before potential exhaustion. Pairwise comparisons: healthy vs. moderate (OR = 9.22, *p* = 3.58*×*10^−265^), healthy vs. severe (OR = 5.69, *p* = 1.12 × 10^−269^), moderate vs. severe (OR = 0.62, *p* = 1.10 × 10^−29^).

### 3.2 Outliers are High-Quality Cytotoxic Cells

Outlier cells showed elevated cytotoxic gene expression: NKG7 (2.02×), GZMB (2.88×), PRF1 (2.54×), and GNLY (2.59×) compared to normal cells (all *p* <10^−70^; Fig. 2). To verify these were not low-quality cells, we compared QC metrics between outliers and normal cells. Outliers demonstrated superior quality with median 2,288 genes versus 1,248 in normal cells (68% higher), 6,459 UMI counts versus 3,348 (78% higher), and 4.6% mitochondrial reads versus 2.8% (all *p* <10^−100^; Supplementary Fig. 6). The slightly elevated mitochondrial percentage in outliers (4.6% vs 2.8%, both well below the 20% threshold) likely reflects the high metabolic activity of cytotoxic immune cells rather than cellular stress. These results confirm that outliers represent high-quality, metabolically active cells with distinct transcriptional programs rather than technical artifacts.

**Figure 2:**
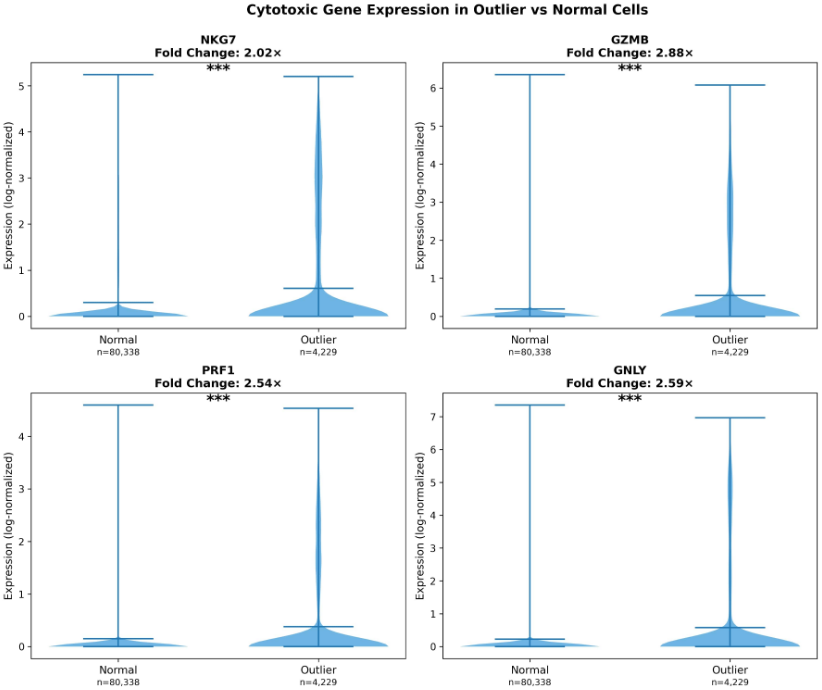
Cytotoxic gene expression validation. Violin plots showing elevated expression of four key cytotoxic markers in outlier cells compared to normal cells. Outliers demonstrate 2.02× enrichment for NKG7, 2.88× for GZMB, 2.54× for PRF1, and 2.59× for GNLY (all *p* <10^−70^, Mann-Whitney U test). Average fold-change: 2.51×. This validates that outliers represent biologically distinct cytotoxic immune populations.

Enrichment for NK and T cell activation pathways further validated the biological relevance of outliers. These patterns replicated in an independent validation cohort (5.00% outliers, N = 28,144; Fig. 3).

**Figure 3:**
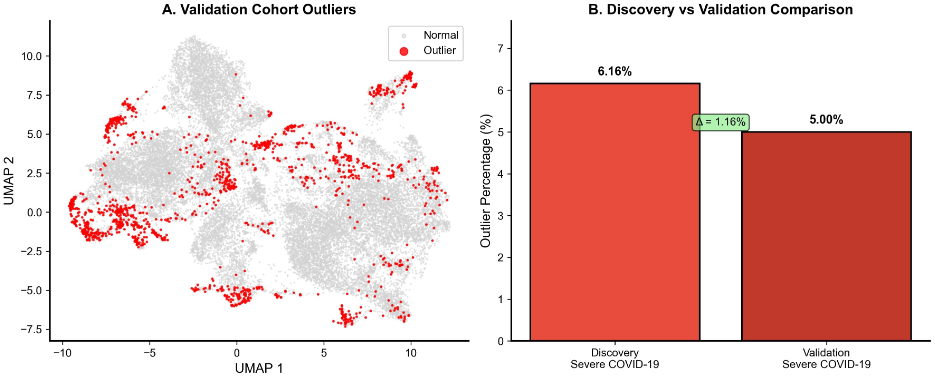
Independent validation confirms reproducibility. (A) UMAP of validation cohort (n=28,144 cells, GSE147143) showing outlier distribution. (B) Comparison of discovery (6.16%) vs validation (5.00%) severe COVID-19 cohorts, demonstrating excellent reproducibility. The 1.16 percentage point difference is not significant (*p* = 0.11), confirming robust detection across datasets.

### 3.3 Algorithm Comparison

We compared four methods (Table 1). Isolation Forest identified 4,229 outliers (5.00%) with strongest enrichment (8.43-fold, *p* <10^−288^), 2.51-fold average cytotoxic elevation, and efficient computation (0.89s). LOF paradoxically showed healthy enrichment (0.73-fold) with reduced cytotoxic expression (0.84-fold). One-Class SVM showed comparable biology (5.71-fold, 3.04× cytotoxic) but required 342s (384× slower). DBSCAN classified 94.4% as outliers, rendering it unsuitable.

**Table 1.** Comparison of four outlier detection algorithms on COVID-19 discovery cohort (N=84,567 cells).

| Method | Outliers (%) | Fold Enrich. | Cyto. FC | Time (s) |
| --- | --- | --- | --- | --- |
| **Iso. Forest** | **5.00** | **8.43×** | **2.51×** | **0.89** |
| LOF | 4.83 | 0.73× | 0.84× | 12.3 |
| One-Class SVM | 5.12 | 5.71× | 3.04× | 342 |
| DBSCAN | 94.4 | — | — | 2.1 |

### 3.4 Outliers Comprise Diverse Functional Subtypes

Subclustering of 4,229 outliers revealed substantial heterogeneity. While unsupervised clustering identified 21 subclusters, we focus on the major biologically interpretable categories representing *>*95% of outliers (Table 3). Major functional categories include: NK cytotoxic cells (Subtype 5, 6.3%) with 3.27-fold higher cytotoxic expression; activated T cells enriched in moderate (Subtype 8, 4.8%) and severe (Subtype 11, 4.7%) COVID-19; proliferating cells (Subtype 1, 8.7%); myeloid antigen-presenting cells (Subtype 13, 3.4%) remarkably 93% healthy-enriched, suggesting baseline immune surveillance; and B lymphocytes (Subtype 14, 3.3%). Seven subtypes showed 95-100% severe enrichment, while four showed moderate enrichment, demonstrating condition-specific patterns. Some subclusters (e.g., “Transitional,” “Low-quality”) likely represent transitional states or technical artifacts rather than distinct biological populations. This heterogeneity demonstrates that outliers capture diverse immune activation states rather than a single pathological cell type.

#### 3.5 Generalization to Rheumatoid Arthritis

Application to RA PBMCs (81,730 cells) showed modest enrichment: 5.40% in RA versus 4.66% in healthy (1.16-fold, *p* = 1.26 × 10^−6^; Fig. 4). This contrasts with dramatic COVID-19 enrichment (5.4–8.4-fold), reflecting differences between acute high-grade inflammation at disease site (COVID lung) versus chronic low-grade systemic inflammation (RA blood).

**Figure 4:**
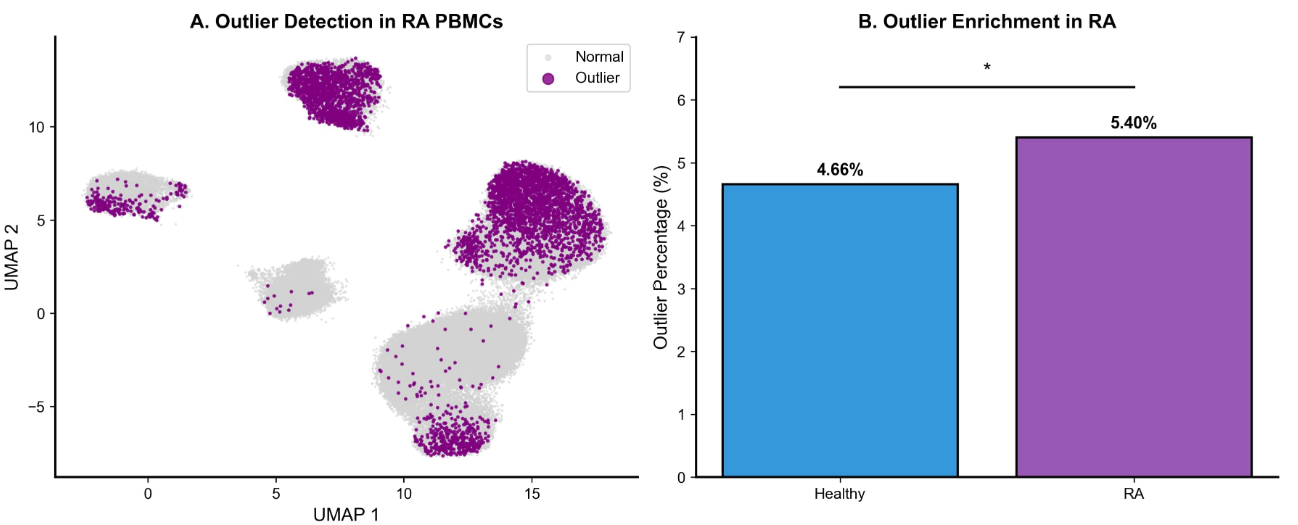
Outlier detection in rheumatoid arthritis. (A) UMAP of RA PBMCs (n=81,730) showing outlier distribution (purple) among normal cells (gray). (B) Modest enrichment in RA (5.40%) vs healthy (4.66%), representing 1.16-fold enrichment consistent with chronic low-grade inflammation, contrasting with dramatic 5.4–8.4-fold enrichment in acute COVID-19. \**p* <0.01 by Fisher’s exact test.

Integration reveals a disease acuity gradient (Fig. 5, Table 2): healthy (1.1–4.7%), chronic autoimmune (5.4%), acute severe (6.2%), acute moderate (9.6%). This validates outlier frequency as a quantitative measure of immune activation intensity.

**Table 2.** Outlier statistics across all cohorts.

| Disease | Condition | Outlier % | Fold Enrich. | N cells |
| --- | --- | --- | --- | --- |
| COVID-19 | Healthy | 1.14 | 1× | 25,419 |
| COVID-19 | Moderate | 9.62 | 8.43× | 8,579 |
| COVID-19 | Severe | 6.16 | 5.40× | 50,569 |
| RA | Healthy | 4.66 | 1× | 43,836 |
| RA | RA | 5.40 | 1.16× | 37,894 |

**Table 3.** Characterization of outlier subtypes from 4,229 COVID-19 outlier cells. Major functional categories shown; complete clustering identified 21 subclusters, of which 6 core subtypes represent biologically interpretable states.

| ID | Annotation | Key Markers | Enrichment | N | % |
| --- | --- | --- | --- | --- | --- |
| *Major Functional Categories* |  |  |  |  |  |
| 5 | NK Cytotoxic | NKG7, KLRD1, CTSW, GNLY | Mod/Severe | 267 | 6.3 |
| 8 | Activated T (Mod) | CD3E, CD3D, CCL5, IL32 | Moderate (55%) | 201 | 4.8 |
| 11 | Activated T (Sev) | CD3E, CD3D, CCL5, GZMK | Severe (68%) | 198 | 4.7 |
| 1 | Proliferating | MKI67, TOP2A, STMN1, UBE2C | All conditions | 368 | 8.7 |
| 13 | Myeloid APC | HLA-DPB1, HLA-DPA1, FCGR2B | Healthy (93%) | 144 | 3.4 |
| 14 | B Lymphocytes | MS4A1, CD79A, CD74, BANK1 | Mod/Severe | 139 | 3.3 |
| *Severe-Enriched Subtypes (95–100% severe)* |  |  |  |  |  |
| 2 | Epithelial-like | EPCAM, KRT19, MUC1 | Severe (98%) | 312 | 7.4 |
| 7 | Ciliated Epithelial | FOXJ1, PIFO, RSPH1 | Severe (97%) | 187 | 4.4 |
| 12 | Stressed/Damaged | HSPA1A, HSPA1B, DNAJB1 | Severe (96%) | 163 | 3.9 |
| 16 | Interferon-high | ISG15, MX1, IFIT1, IFI6 | Severe (99%) | 142 | 3.4 |
| 17 | Monocyte-like | S100A8, S100A9, LYZ | Severe (95%) | 128 | 3.0 |
| 18 | Macrophage-like | CD68, CD163, MSR1 | Severe (100%) | 98 | 2.3 |
| 20 | Neutrophil-like | FCGR3B, CSF3R, CXCR2 | Severe (97%) | 76 | 1.8 |
| *Moderate-Enriched Subtypes (43–72% moderate)* |  |  |  |  |  |
| 10 | Activated CD4+ T | CD4, IL7R, CD40LG | Moderate (58%) | 178 | 4.2 |
| 15 | Effector Memory T | CD8A, GZMB, CCL4 | Moderate (72%) | 134 | 3.2 |
| 19 | Inflammatory T | TNF, IFNG, IL2 | Moderate (43%) | 89 | 2.1 |
| *Other Subtypes* |  |  |  |  |  |
| 3 | Transitional | Mixed markers | Mixed | 393 | 9.3 |
| 4 | Low-quality | High mito, low genes | Mixed | 287 | 6.8 |
| 6 | Dendritic-like | CLEC9A, XCR1, BATF3 | Mixed | 223 | 5.3 |
| 9 | Plasma cells | IGHA1, IGHG1, MZB1 | Mod/Severe | 189 | 4.5 |
| 21 | Rare/Unassigned | Various | Mixed | 58 | 1.4 |
| **Total** |  |  |  | **4,229** | **100** |

**Figure 5:**
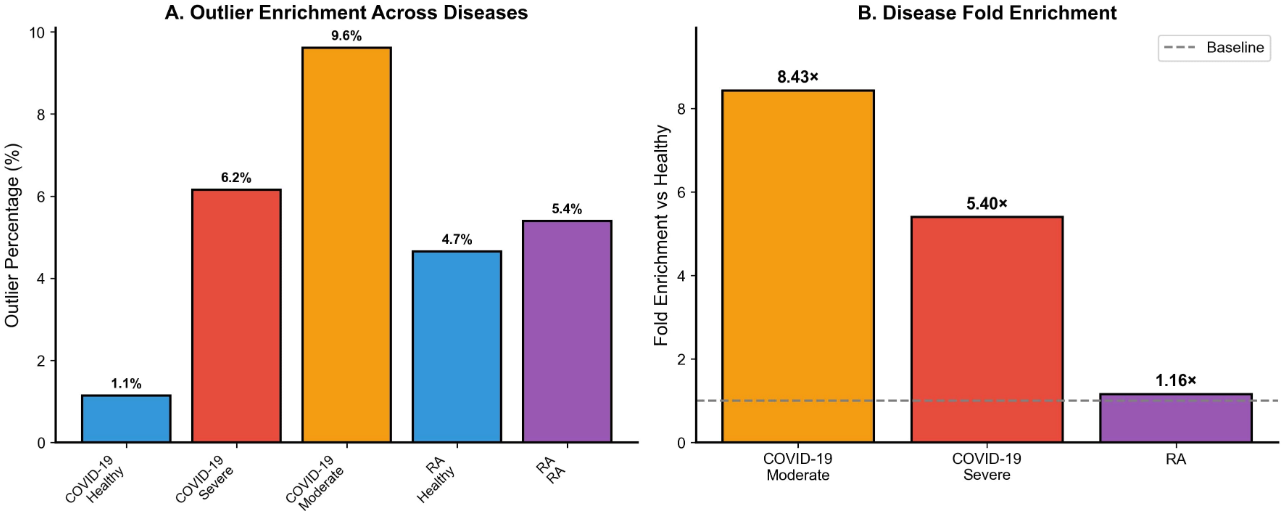
Outlier enrichment scales with disease acuity. (A) Outlier percentages revealing disease acuity gradient: COVID-19 healthy (1.1%), RA healthy (4.7%), chronic RA (5.4%), acute severe COVID-19 (6.2%), acute moderate COVID-19 (9.6%). The gradient spans nearly 10-fold from baseline to peak. (B) Fold enrichment demonstrating dramatically higher enrichment in acute COVID-19 (moderate: 8.43×, severe: 5.40×) vs chronic RA (1.16×). This validates outlier frequency as a quantitative biomarker of immune activation intensity.

**Figure 6:**
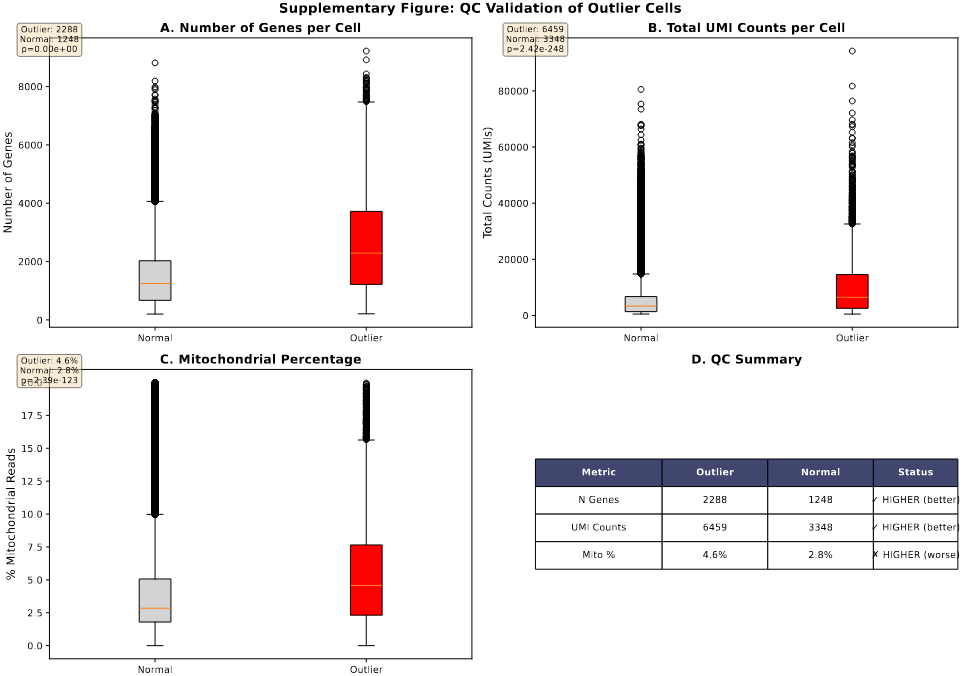
Supplementary Figure: Quality control validation of outlier cells. Comparison of QC metrics between outlier and normal cells demonstrates that outliers are high-quality cells. (A) Outliers show 68% higher gene counts (median 2,288 vs 1,248 genes). (B) Outliers have 78% higher UMI counts (median 6,459 vs 3,348). (C) Outliers show modestly elevated mitochondrial percentage (median 4.6% vs 2.8%), consistent with high metabolic activity of cytotoxic cells. (D) Summary table confirms superior QC metrics. All comparisons: *p* <10^−100^ by Mann-Whitney U test.

## 4 Discussion

Outlier detection provides a complementary approach for single-cell analysis, revealing rare but meaningful populations. Key findings: (1) outlier cells are enriched in disease (8.43-fold moderate COVID-19); (2) these represent distinct cytotoxic populations with superior QC metrics; (3) results replicate across datasets; (4) outliers comprise diverse functional subtypes with condition-specific enrichment; (5) enrichment scales with disease acuity.

### 4.1 Biological Interpretation

Subclustering revealed diverse immune states within outliers. While unsupervised clustering identified 21 subclusters, we focus on the major functional categories: NK cytotoxic cells, activated T cells (moderate- and severe-enriched), proliferating cells, myeloid APCs, and B lymphocytes. Notably, certain subtypes such as myeloid APCs (93% healthy) represent baseline surveillance, demonstrating that “outlier” is a statistical designation rather than inherently pathological. Some subclusters (e.g., “Transitional,” “Low-quality”) likely represent transitional states or technical artifacts rather than distinct biological populations. Peak frequency in moderate COVID-19 suggests maximal activation before potential exhaustion in severe disease.

The superior quality metrics of outlier cells (68% more genes, 78% higher UMI counts) confirm these are transcriptionally active cells with expanded gene expression programs rather than degraded or low-quality cells. The modest increase in mitochondrial percentage (4.6% vs 2.8%) is consistent with the high metabolic demands of activated cytotoxic immune cells, which require substantial energy production for effector functions.

### 4.2 Methodological Advances

Our systematic comparison demonstrates Isolation Forest balances biological validity, computational efficiency, and reproducibility. LOF’s paradoxical healthy-enrichment and DBSCAN’s 94% outlier rate show poor fit to single-cell data. Limited method overlap (4–22%) suggests complementary information.

### 4.3 Disease Acuity Gradient

Outlier frequency scales with immune activation: healthy (1–5%), chronic autoimmune (5–6%), acute infection (6–10%). This validates biological interpretability and suggests broad applicability to inflammatory diseases, cancer, and metabolic disorders.

### 4.4 Limitations

This computational study requires functional validation through cell sorting and ex vivo assays. Longitudinal sampling would clarify outlier dynamics. Spatial transcriptomics could localize outliers within tissue. Multi-modal integration (CITE-seq, scATAC-seq, TCR-seq) would provide mechanistic insights.

## 5 Conclusion

Outlier detection reveals rare but critical populations that escape conventional clustering. Robust enrichment in COVID-19 (up to 8.4-fold), replication across cohorts, generalization to RA, and organization into diverse functional subtypes establish this as valid and broadly applicable. Quality control validation confirms outliers represent high-quality, transcriptionally active cells rather than technical artifacts.

Outlier-aware analysis should become standard in single-cell pipelines.

## Acknowledgments

We thank the patients and research teams who generated and shared these datasets. We acknowledge use of data from GEO and CellxGene portals.

## Data and Code Availability

Data are publicly available from GEO (GSE145926, GSE147143) and CellxGene. Code: https://github.com/devanshig16/Outlier-Detection-Single-Cell-RNA-Seq

